# An accessible and versatile deep learning-based sleep stage classifier

**DOI:** 10.1101/2022.08.24.504155

**Authors:** Jevri Hanna, Agnes Flöel

## Abstract

Manual sleep analysis for research purposes and for the diagnosis of sleep disorders is labor-intensive and often produces unreliable results, which has motivated many attempts to design automatic sleep stage classifiers. With the recent introduction of large, publicly available hand-scored polysomnographic data, and concomitant advances in machine learning methods to solve complex classification problems with supervised learning, the problem has received new attention, and a number of new classifiers that provide excellent accuracy. Most of these however have non-trivial barriers to use. We introduce the Greifswald Sleep Stage Classifier (GSSC), which is free, open source, and can be relatively easily installed and used on any moderately powered computer. In addition, the GSSC has been trained to perform well on a large variety of electrode set-ups, allowing high performance sleep staging with portable systems. The GSSC can also be readily integrated into brain-computer interfaces for real-time inference. These innovations were achieved while simultaneously reaching a level of accuracy equal to, or exceeding, recent state of the art classifiers and human experts, making the GSSC an excellent choice for researchers in need of reliable, automatic sleep staging.

## Introduction

Analysis of sleep stages for diagnosis of various sleep disorders, as well as analysis on more sophisticated microstructure of sleep like slow oscillations, spindles, and their coupling for research purposes (Rasch and Born 2013 for review), has become an important goal in clinical and research context. Sleep consists of a rich diversity of neural and physiological stages, which typically unfold in semi-regular cycles throughout the sleep period. These stages have distinct signatures that can be measured with polysomnography (PSG), which includes the measure of neurophysiology with electroencephalogram (EEG), as well as ocular (EOG) and muscular (EMG) activity. Established guidelines (Rechtschaffen, 1968, Silber et al. 2007) allow for manual classification of PSG data into discrete sleep stages in 30 second increments, but this is a highly laborious process, requiring as much as two hours to classify a single night’s sleep, even for a trained expert (Vallat and Walker, 2021). In addition to the costs of manual classification, there is also substantial disagreement between expert sleep stage scorers (70-80% agreement) and even *within* the same expert scorer at different times (90% agreement) (Rosenberg and Van Hout, 2013, Younes et al, 2016, Muto et al, 2018), which introduces a non-trivial degree of unreliability to both research findings and clinical diagnosis based on manual sleep staging. This variability is an inevitable consequence of the relative indeterminacy involved in applying sleep stage criteria to highly complex and variable human polysomnographic data. Sleep scorers make decisions on the basis of e.g. occipital alpha for a certain period of time, sleep spindles, K-complexes, certain types of eye movements, or overall EEG amplitude, to name just a few. On top of this, there are extensive contextual rules that specify under what circumstances one stage can follow another, adding a further layer of complexity and subjectivity. There is self-evidently wide interpretative discretion among sleep scorers as to how to identify these phenomena and how exactly to balance these rules against each other. The same indeterminacy also makes algorithmic sleep staging with traditional, analytical approaches difficult. Nevertheless, there have been many attempts, dating back to at least the 1990s, though these generally have not demonstrated robust generalizability (see Sun et al. 2017 for references and discussion).

Meanwhile however, in the previous decade, tens of thousands of hours of expert-scored PSGs have become publicly available (Zhang et al., 2018, sleepdata.org), and an explosion of algorithmic innovation and increased computing power has opened the possibility to train machine learning and deep learning models on these large datasets. Given the costliness and unreliability of manual scoring on the one hand, and the limitations of analytical approaches on the other hand, it is unsurprising that automatic sleep staging with machine/deep learning immediately became a field of intense focus and development. Deep learning in particular has proven to be very well suited to solving many highly complex pattern recognition problems that had not been satisfactorily solved with prior methods (LeCun et al, 2015). In a short period of time, a large number of classifiers have been developed utilising machine learning or deep learning to successfully score sleep stages (for review see Fiorillo et al, 2019). However, even though this has produced a significant improvement to the state of the art, there are still several drawbacks to most currently available sleep classifiers. As Vallet and Walker (2021) point out in the introduction of their own sleep classifier, YASA, these classifiers have barriers which make them less accessible, such as either 1) costing money or requiring expensive software to run (e.g. MATLAB), 2) requiring more technical knowledge to configure and use than many researchers have at their disposal, or 3) requiring data transmission to an external server.

To remedy these shortcomings and provide high-quality, automatic sleep staging to a larger community of researchers, we introduce here the Greifswald Sleep Stage Classifier (GSSC), with the following overall goals in mind: First, as with YASA (Vallet and Walker, 2021), we have endeavoured to produce a sleep stage classifier that is open source, freely available and not dependent on paid software, relatively easy to install and use, and can be run locally on any reasonably powered PC. Second, the GSSC was trained in order to achieve high performance also on less common electrode arrangements, including EOG only. Third, the GSSC has been designed such that it can be straightforwardly integrated into brain computer interfaces or closed loop brain stimulation systems, with minimal processing overhead. Fourth, we sought to make improvements to the overall accuracy of the classifier in relation to the state of the art.

## Methods

### Architecture

The GSSC uses a relatively simple architecture that requires minimal preprocessing or assumptions about relevant data features. The first part is a signal processing module that uses Resnets to convert the signal(s) into an abstract representation - expressed as a vector of size 1280. Resnets are a form of convolutional neural network that utilize skipping connections to alleviate some of the typical problems encountered with deeper networks (He et al, 2016, Wu et al, 2019). Two separate Resnets were trained; one for EEG and one for EOG. During prototyping, we found that the EOG Resnet does not benefit from significant depth, whereas the EEG Resnet improves significantly from added layers; here we added four Resnet blocks with each downsampling block. The next stage is a mixing and compression network, which - in the case of either just one EEG or just one EOG channel - compresses the vector of size 1280 into a vector of size 512, and in the case of using both EEG and EOG channels, mixes and compresses the EEG and EOG vectors (2×1280) into a vector of 512. Finally, this compressed vector of size 512 is either sent to a three-layer fully-connected network that decides the sleep stage, or passed onto a bidirectional Gated Recurrent Network (GRU) (Cho et al. 2014), which decides the sleep stage based on both the compressed vector and a hidden state which encodes the preceding/subsequent context. Network architecture is depicted schematically in fig. 1.

**Figure 1:**
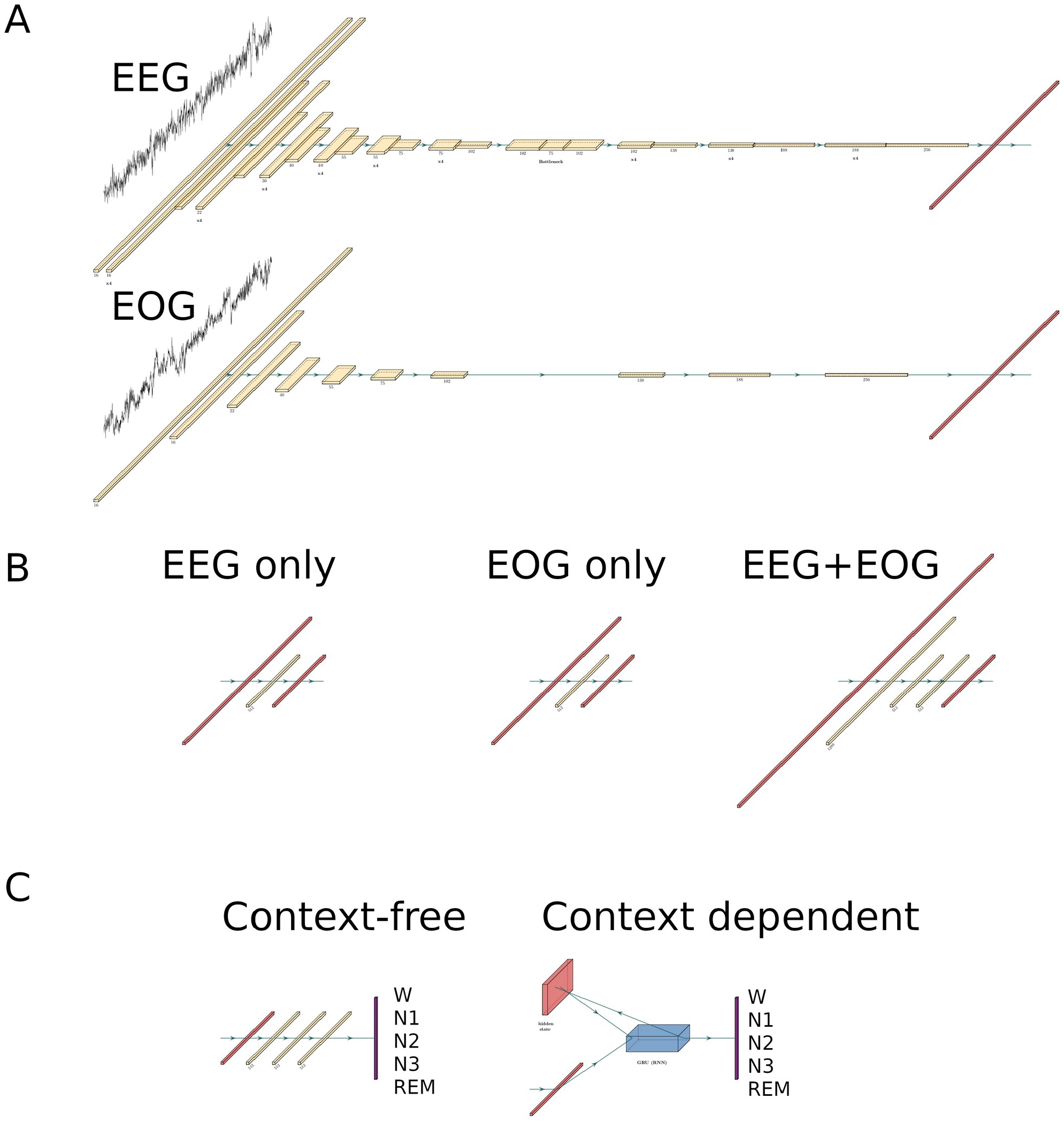
Neural network architecture. **A)** EEG and/or EOG input are processed by one-dimensional convolutional Resnets that progressively reduce the extent of the input in signal space while increasing it in abstract, feature space. The final output of the EEG or EOG net is flattened into a vector of length 1280. **B)** This vector is compressed by a one layer fully-connected network into a vector of length 512. If both EEG and EOG are used, then the concatenated vectors from A are mixed and compressed by a three layer fully-connected network into a vector of length 512. **C)** The compressed vector is then fed into either a three layer fully connected network that outputs a vector of length five that encodes the inferred sleep stage; this is context-free inference. Context dependent inference feeds the compressed vector along with a hidden state into a Gated Recurrent Unit (GRU). The GRU produces an output vector of length five as well as a new hidden state, which is then fed back into the GRU for the next sleep stage.

### Data sets

We briefly list here the datasets and how they were implemented. For more information see sleepdata.org (Zhang et al, 2018). All datasets used the AASM system for sleep scoring (Silber et al. 2007), except the Sleep Health Heart Study (SHHS), which used Rechtschaffen and Kales (Rechtschaffen, 1968); the latter was rescored to be compatible with the other datasets.

### Sleep Health Heart Study 2 (SHHS2)

The SHHS (Quan et al. 1997) is a large set of home PSGs assembled through 5 cohorts throughout the United States, primarily for the purpose of researching the connection between sleep-related breathing and cardiovascular disease. All participants were at least 40 years of age. We restricted ourselves here to the second phase of the project SHHS2, collected between 2001-2003, and used here a sample of 936 PSGs from the total 3,295 PSGs. Relevant electrodes include C3-A2, C4-A1, and left and right EOG. This study used the Rechtschaffen and Kales system, which we rescored to the AASM system (all N4 stages become N3).

### Cleaveland Family Study (CFS)

The CFS (Redline et al, 1995) is a longitudinal (1990-2006) study focussing on sleep apnea in the United States. It features a particularly high proportion (46%) of Black American participants. We use here all 730 of the available PSGs. Relevant electrodes include C3-Fpz, C4-Fpz, and left and right EOG.

### Nationwide Children’s Hospital Sleep DataBank (NCHSDB)

The NCHSDB (Lee et al, 2021) is a large pediatric dataset of PSGs collected in the United States, with most participants under the age of 10. We use here a sample of 665 of the 3,984 PSGs.

Relevant electrodes include F3-M2, F4-M1, C3-M2, C4-M1, and left and right EOG.

### Wisconsin Sleep Cohort (WSC)

The WSC is a still ongoing longitudinal study focussing on various sleep disorders, collected in the United States. We use here a sample of 983 PSGs from the second stage of the project, collected with the Grass Comet Lab system (2009-present). Relevant electrodes include F3-M2, C3-M2, O1-M2, and left and right EOG. For more information see https://sleepdata.org/datasets/wsc and Young et al. (2009).

### Training, testing, and validation partitions

A combined total of 3,314 nights of manually scored PSGs, comprising 29,299 hours and 3,515,889 individual 30s epochs, derived from the four datasets listed above were used for training, testing and validating the networks. Additionally, the DREEM dataset (Guillot et al, 2020) was used as a final validation and point of direct comparison with a few of the most recent other classifiers. 80% (2652 PSGs) of the full dataset were used for training. 10% (331) were used for testing (model prototyping, hyperparameter selection, assessment of overfitting), and a final 10% were used for validation; performance on these 10% as well as on the DREEM dataset are the basis for all reported results.

### Preprocessing

All PSGs were finite impulse response filtered with a bandpass of 0.3-30Hz. This band captures the most relevant oscillatory phenomena in human sleep, and safely excludes all line noise (50/60Hz). Signals were Z-transformed on a per-channel, per-PSG basis. The right EOG channel was subtracted from the left EOG channels to form a single HEOG channel. All signals downsampled to 85.33 Hz, which reduces a 30s section to 2560 samples, the length of signal input to the networks. Data were otherwise not cleaned or pruned in any way.

### Signal permutations and data augmentation

Every PSG in the training partition was trained multiple times, each time with a different signal permutation. Possible permutations include 1) an EEG channel, 2) the HEOG channel, 3) an EEG channel and HEOG channel together. Each possible EEG channel in a dataset was the basis of a permutation under conditions (1) and (3). In addition to these permutations, signals could also be flipped in polarity, i.e. by multiplying them by −1. This would mean for example that a dataset with two EEG channels and one EOG channel would have 14 possible permutations. The motivation for polarity flipping is to approximate the intrinsic relativity of EEG signals; every signal can easily change polarity simply by changing the EEG reference. Training under bipolarity, in addition to significantly augmenting the dataset size, has the goal of forcing the classifier to learn more abstract properties of the signal, resulting in a more flexible classifier that is likely to perform well under a larger variety of PSG recording set-ups and reference channels. Earlier prototypes also made use of the chin EMG channel, but did not noticeably improve performance, similar to what was reported in Perslev et al. (2021). In the YASA classifier (Vallat and Walker, 2021), only one of the top 20 classification features was EMG based, ranking at 18th. This suggests that chin EMG contributes relatively little unique information to sleep stage classification.

### Training procedure

Neural networks were implemented within the PyTorch framework (v.1.10.2). For 20 training epochs, the 2652 training PSGs were cycled through in random order. In order to fit within GPU memory constraints, PSGs were divided into 128 batches each with a roughly equal amount of contiguous, 30s sections. A forward and backward pass was calculated on each batch, moving sequentially through the PSG. The forward and backward pass had two modes: context-free and context-aware. Both modes shared a common path for the first two stages, namely the signal processing and mixing/compression modules (see Architecture, Fig. 1 A/B). After this point, they branched off, with the compressed vector going to the fully-connected, three-layer decision network in the case of context-free mode, which produced a one-hot vector of length 5, encoding for five possible sleep stages. This vector was compared against the correct sleep stage with a cross entropy loss function, and this loss was back-propagated through the decision network, mixing/compression, and signal processing modules. In the case of context-aware mode, the compressed vector and the previous hidden state was sent to the GRU network, which outputs a one-hot vector of length 5, again encoding for five possible sleep stages, and a new hidden state for the next batch. The loss was calculated in the same way as in context-free mode, and back-propagated through the GRU network, mixing/compression, and signal processing modules. After the losses had been back-propagated in both modes, the weights were updated, and the process was repeated for the next batch. For the first 30s section of a PSG, the initial hidden state for the GRU network was set to all zeros. Because the models were trained in both modes simultaneously, the signal processing and mixing/compression modules learned to produce representations which could be used interchangeably with either the context-free decision network, or the context-aware GRU network. Weights were updated with the AdamW optimizer (Zhang Z., 2018) with hyperparameters beta1=0.9, beta2=0.999, and learning rate=3e-5. Dropout (Hinton et al. 2012) was applied after every layer during training at the rate of 0.5. Because the frequency of sleep stages is severely imbalanced, it is necessary to provide the loss function with class weights to prevent the model from simply learning to guess blindly according to sleep stage probability. We adopted the weights used for the YASA algorithm (Vallat and Walker, 2021): Wake: 1, N1: 2.4, N2: 1, N3: 1.2, REM: 1.4, changing only the N1 weight slightly from 2.2 to 2.4 on the basis of prototyping.

### Assessment measures

As a primary measure of performance, we use the Matthews Correlation Coefficient (MCC), which requires high true positives and negatives as well as low false positives and negatives to produce a good score, and is particularly well-suited for evaluating results on unbalanced datasets (Chicco and Jurman, 2020). In the interest of comparability with other studies as well as offering quick, intuitive results, we also report here F1 for each sleep stage, F1 macro, simple accuracy, Cohen’s Kappa, and confusion matrices (see Menghini et al. 2021 for discussion of these). All of these metrics are calculated for each PSG separately. Medians across all PSGs rather than means are reported to prevent distortion from outlier PSGs.

### Permutation consensus

The classifier can infer sleep stage from any combination of EEG and EOG channels, or from only one of the two. It is therefore possible to make multiple inferences from the same PSG using different channel combinations. It also possible to calculate the certainty of that inference by calculating the cross entropy of the log softmax vector of 5 that is output by the classifier against the inferred sleep stage. Results indicated that inferences with high certainty also tended to be correct more often than lower certainty inferences (fig. S1). By adopting the inference of the permutation with the highest certainty, we can increase the accuracy of the classifier.

### Selection of optimal network and validation

Throughout training, the performance of the classifier on training data continually improves. This does not necessarily indicate however that the final state of the network is the best one; overfitting on the training set can occur. To ascertain the optimal stopping point for training, we assessed classifier performance at the end of each training epoch on the testing dataset (331 PSGs), and identified the point at which performance peaked. This point was chosen as the optimal network which would be used in the final, validation step.

As a final confirmation of the performance of the model on unseen data, we assessed optimal network classifier performance on the validation dataset (331 PSGs). In addition to this, we also assessed performance on the DREEM dataset, both on healthy participants (n=25) and those with sleep related breathing disorders (n=55). The DREEM dataset was rated by five expert sleep scorers, and we assessed the classifier against the majority consensus of their scoring. In parallel to this, we assessed the YASA algorithm (Vallat and Walker, 2021) on both the validation dataset and the DREEM dataset for a direct comparison of the two. Finally, for the DREEM dataset only, we report the performance of two other state of the art classifiers, U-Sleep (Perslev et al, 2021) and that of Stephansen et al (2018), using data kindly provided to us by Raphael Vallat (Vallat and Walker, 2021). In order to assess the flexibility of the GSSC and YASA classifiers, we assessed performance on all possible combinations of EEG/EOG/EMG channels. EMG combinations were implemented for YASA only, as GSSC does not make use of EMG channels, and combinations without EEG were implemented for GSSC only, as YASA requires the use of an EEG channel. Also, for the GSSC classifier we made use of the consensus of permutations assessment (see assessment measures above).

## Results

### Training and model selection

During training, loss steadily declined and accuracy steadily increased; by the 20th training epoch these were near asymptote (fig S2). Performance on the test set however oscillated up and down across the 20 training epochs, and did not reach convergence (see fig S3). For this we reason, we selected three epochs that had both a high Matthews Correlation Coefficient and a high F1 Macro score. The reason for balancing across these two measures is that a high F1 Macro ensures that accuracy for all five sleep stages is relatively good; due to the severely unbalanced nature of sleep stages, a model could perform poorly in a less common sleep stage but still have a high overall MCC or accuracy. The weights from these three training epochs were averaged and the resulting weights were then used to infer sleep stages on the validation datasets. Weight averaging across multiple epochs throughout training has been shown to improve accuracy and generalisation to unseen data (Izmailov et al. 2018).

### Context-aware and context-free inference

To assess the influence that contextual information has on inference accuracy throughout the PSG, we calculated accuracy by PSG epoch across the 355 PSGs from the validation set (including DREEM) that had at least 875 30s epochs (7.5 hours). This was done under three contextual conditions: 1) the optimal, bidirectional inference - where for a given PSG epoch both the preceding and following context were taken into account - was used as the baseline. This type of inference would be used in offline PSG scoring 2) In the forward inference, only the preceding context could be used for inference. This type of inference would be used in e.g. real-time/BCI inference. 3) Context free inference uses no context at all. Results are shown in fig. 2. These indicate that forward inference begins at around a 2% accuracy disadvantage relative to bidirectional inference, that linearly decreases to about 1% over the course of the PSG. Context free inference begins at a disadvantage somewhat below 4% relative to bidirectional inference, which sharply decreases to around an 8% disadvantage by around 2 hours, then declines more slowly until around 5 hours, after which the relative disadvantage seems to asymptote. These results underline the critical role of context in accurate sleep stage inference.

**Figure 2:**
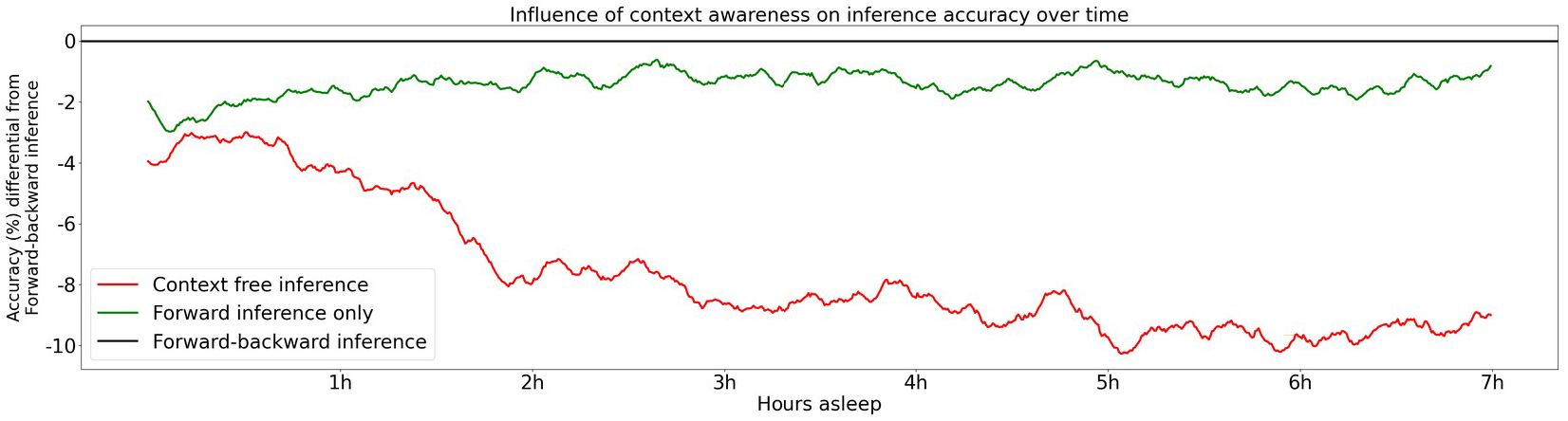
Contribution of contextual information to accuracy. The model performs optimally when making use of both preceding and subsequent epochs (forward-backward inference) - this optimal performance is the baseline at 0. The red and green lines indicate the loss in accuracy for context free inference and forward inference (preceding context only), respectively, across time for 355 PSGs in the validation set.

### Performance on DREEM dataset in relation to other classifiers

Accuracy, F1 Macro, MCC, and Cohen’s Kappa scores for GSSC and three other recent classifers are given in fig 3. Table S1 shows the exact numbers, and fig SX shows the F1 scores for individual sleep stages. These indicate an accuracy advantage of 0.7% for Perslev et al (2021) over GSSC on the Healthy cohort (n=25), and an accuracy advantage of 2% for the GSSC over Perslev et al. (2021) on the Obstructive cohort (n=55). In general, performance seems to be as good or better than Perslev et al. (2021). Because the latter is more accurate than YASA and Stephansen et al. (2018), this indicates that GSSC performs at the current state of the art, and offers the highest possible accuracy among the classifiers compared here.

**Figure 3:**
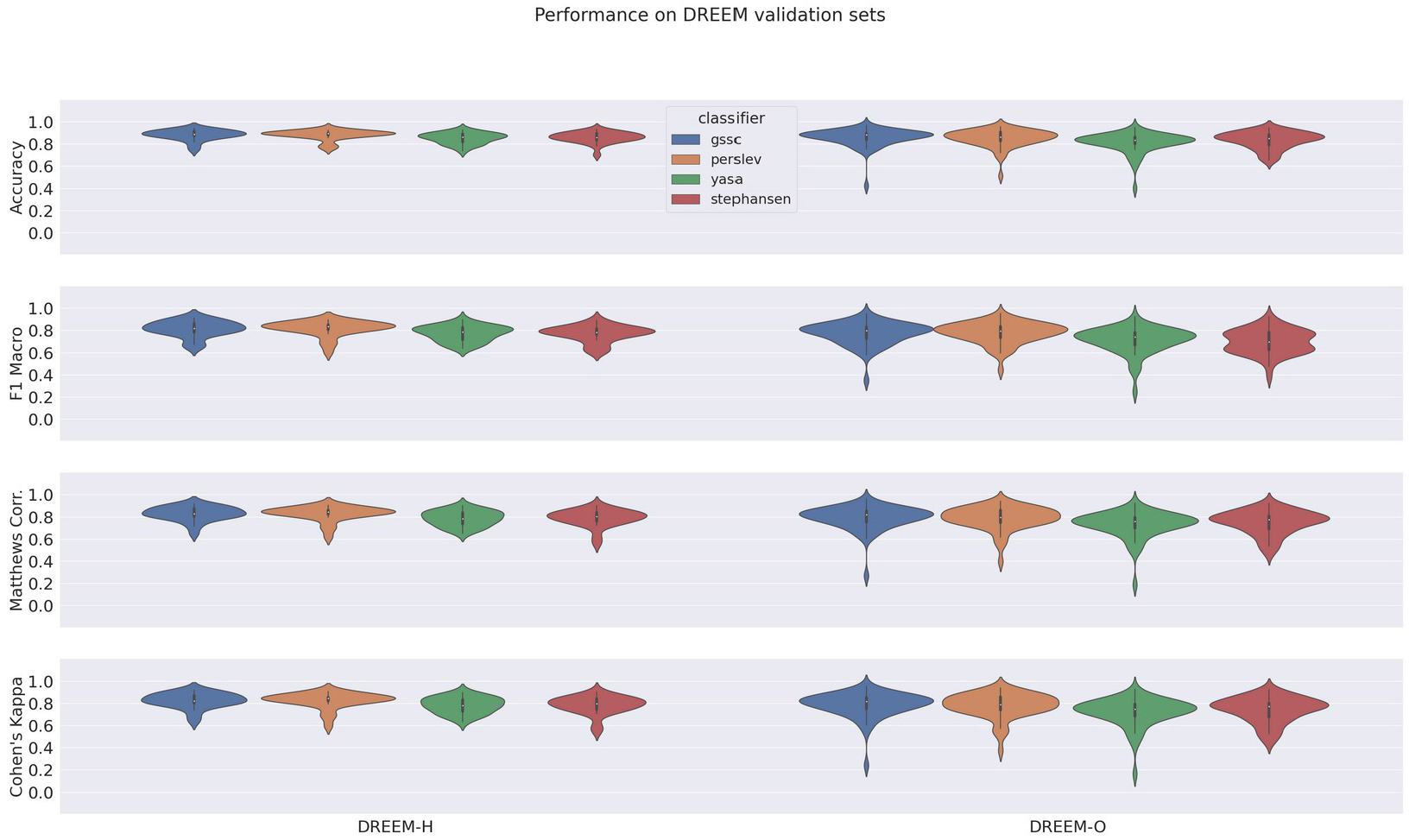
Violin plots of performance over DREEM dataset for four classifiers. Measures shown here include Accuracy, F1 Macro, and Matthews Correlation Coeffecient.

Confusion matrices in figure 4 show that, comparable to other classifiers, the main errors were confusing N1 for N2, and to a much lesser extent confusing N1 for Wake, N2 for N3, and vice versa. This pattern of confusion is consistent with that of both expert human raters and other automatic classifiers, and, aside from the expected, relatively poor performance on N1, shows high accuracy for all sleep stages.

**Figure 4:**
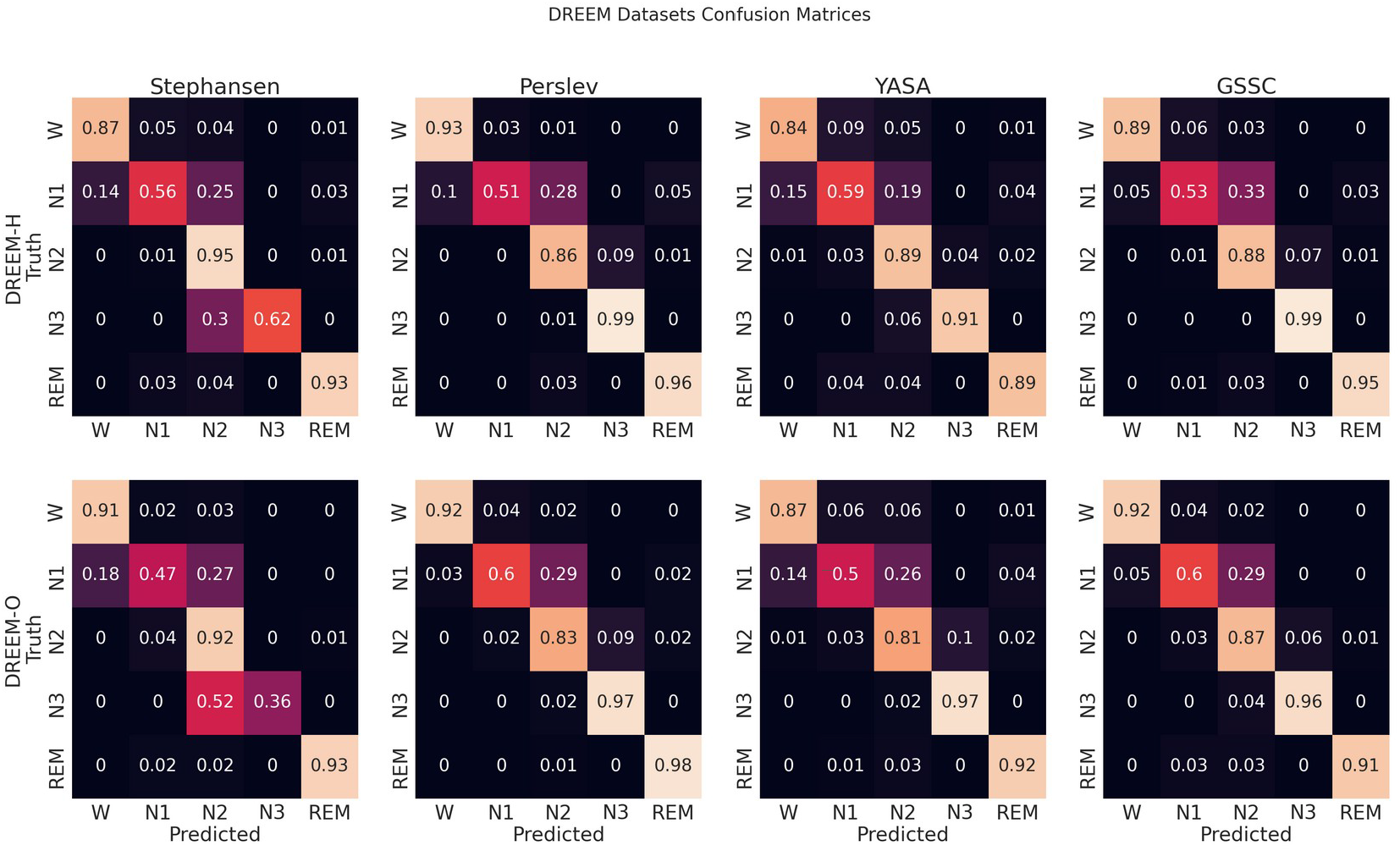
Confusion matrices. These row-normalised confusion matrices show the inferential behaviour for four, recently developed, high performance classifiers, including GSSC. The diagonal indicates the accuracy, and off-diagonal elements show how the true sleep states tended to be misclassified. The first row shows results for the DREEM Healthy dataset (n=25), and the second for the DREEM Obstructed dataset (n=55).

The violin plots in figure 5 show the performance across all validation sets for GSSC and YASA. Mean accuracy is well over 80% in all cases, and outperforms YASA by at least a few points across all datasets. Variance across PSGs also tends to be small except with CFS, NCHSDB, and DREEM-O; these three datasets were either composed of children or pathological populations. Figure 6 shows validation performance for different channels in isolation. All EEG channels perform above 80% mean accuracy. Left and right EOG as well as the difference of the two (HEOG) all have over 80% mean accuracy. Variance in performance across channels for GSSC and YASA was systematically compared with a liner mixed effects (LME) model that took accuracy for each individual EEG channel in each PSG in the validation set as data points, and estimated the fixed effects of channel and classifier on accuracy, with MCC performance on channel C3 as a baseline condition. Estimated effects with their confidence intervals are depicted in fig. 6. These show that for the GSSC, the performance of the different channels tended to cluster within a few percent above or below the baseline. For YASA on the other hand, only channel C4 performed comparably to their recommended C3 channel, and the others tended to be about 5% less accurate. This demonstrates the superior versatility of the GSSC classifier on diverse EEG channels.

**Figure 5:**
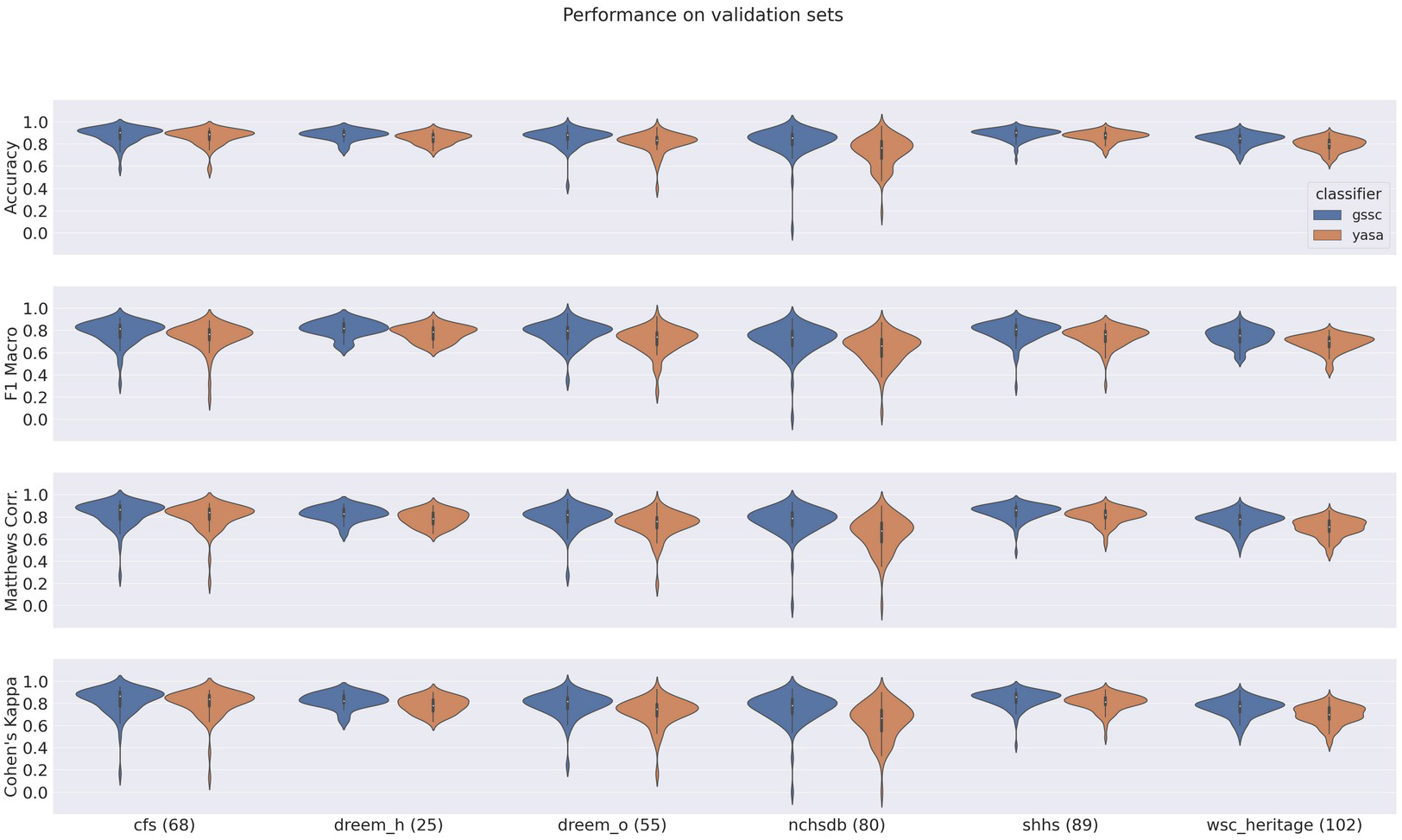
Violin plots of performance over all validation datasets for YASA and GSSC. Measures shown here include Accuracy, F1 Macro, and Matthews Correlation Coeffecient and Cohen’s Kappa. Numbers in parentheses by the dataset name indicate the number of PSGs within that validation set.

**Figure 6:**
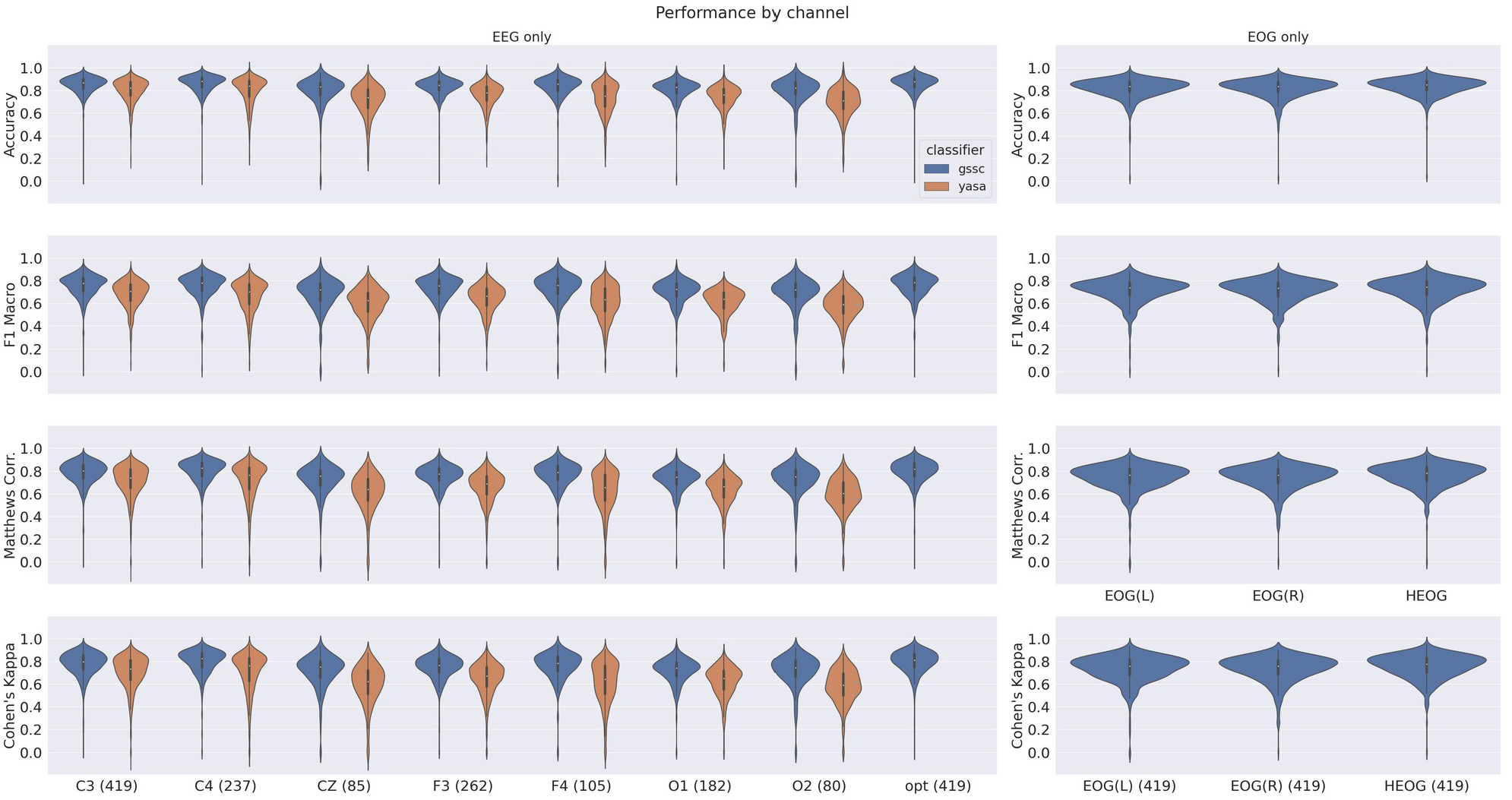
Violin plots for single-channel performance on the validation datasets for YASA and GSSC. Measures shown here include Accuracy, F1 Macro, and Matthews Correlation Coeffecient and Cohen’s Kappa. Numbers in parentheses by the dataset name indicate the number of PSGs which had that channel available.

**Figure 7:**
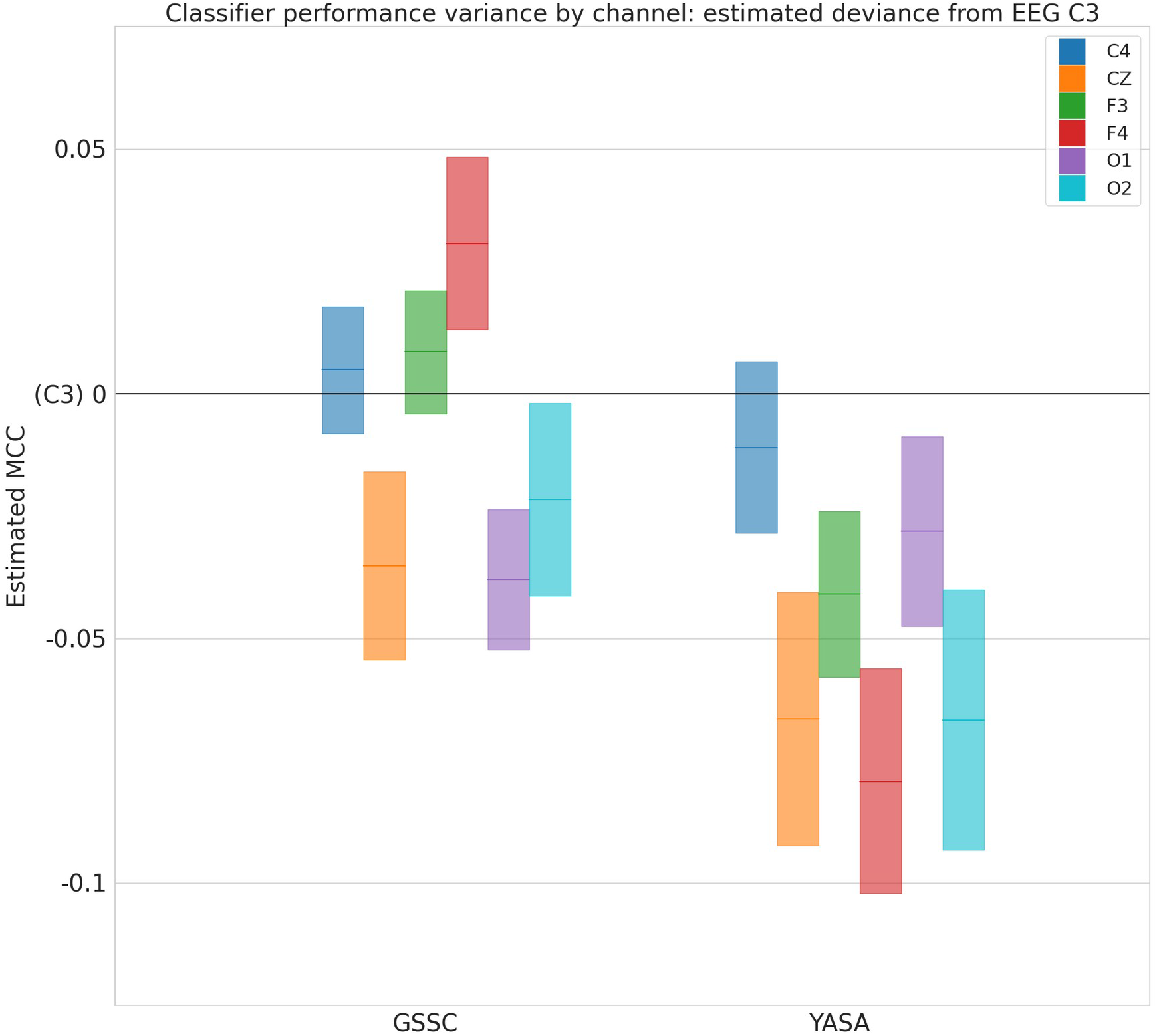
Performance variance across channels. Deviance in Matthews Correlation Coeffecient for different EEG channels and GSSC and YASA classifiers from baseline of EEG C3, as estimated by a linear mixed effects model. Blocks show 95% confidence intervals around the estimated deviance (solid, colored lines).

## Discussion

We have developed an automatic sleep stage classifier that is free and requires no paid software, is easy to install and use, and can be run locally on a moderately well-powered PC. The YASA classifier (Vallat and Walker, 2021) also has these properties, but we have added features to our classifier that will make it preferable for many cases, including greater versatility, easy integration into brain computer interfaces (BCI), and overall improved accuracy. We discuss each of these in turn.

### Free availability and easy access

There has been rapid progress in the sophistication of automated sleep staging technologies, but it has not always been a priority to make this accessible. We developed GSSC within Python, which is free and can be run on any operating system, including Linux, which is also free, meaning that the GSSC can be run entirely on free software if desired. This offers high quality, automatic sleep staging to a much broader base of users who, either because of legal or resource constraints, are unable to use most of the other previously developed classifiers. Even researchers who do have the resources for paid software may nevertheless prefer to work within the free, open-source ecosystem for any number of reasons, not least of which because of the frictions that paid software often impose on their usage with licensing. The GSSC is at present built to work with MNE-Python (Gramfort et al. 2013, website:mne.tools), an M/EEG analysis Python package that is also free, open source, and community developed, though it could easily be adapted to work with any number of EEG analysis programs.

### Versatility

The training strategy we used has produced a versatile classifier that performs well with a broad array of possible electrode configurations. This allows the use of fewer EEG electrodes during recording, or to seamlessly move to backup channels if the more standard channels fail during a recording. GSSC is the only classifier of the four compared here to allow inference with only a single EOG channel, and furthermore with excellent accuracy (>80%). This is concordant with the YASA classifier, which found that EOG absolute power was the single most important feature for sleep stage classification out of all the apriori defined features of the EEG/EOG/EMG signals that they used (Vallat and Walker, 2021). Highly accurate EOG-only inference is of particular use for more portable, 1-2 channel miniature systems designed for home testing (Gao et al. 2021).

### Brain-Computer Interfaces

Recent advances in the relevant hardware and software have fostered increased interest and development in brain-computer interfaces (BCI) (Reza et al. 2019 for recent review). One special case of BCI that is of particular relevance here is the use of closed-loop stimulation, whereby some form of stimulation is given to the participant on the basis of their PSG/EEG activity; such systems have recently been successfully applied during sleep with the goal of modulating Slow Oscillations (e.g. Besedovsky et al. 2017, Kristoffer et al. 2021). For such systems targeting the enhancement of sleep oscillations, it is extremely useful to be able to assess the current sleep stage of the participant. GSSC is relatively easy to integrate into BCIs/closed-loop systems. This functionality is made possible in part by using a recurrent neural network for context awareness. Like human sleep stage scorers, all of the above cited classifiers make use of sleep stage context when performing inference. The GSSC implements context-awareness by use of a Gated Recurrent Network, which takes a hidden state as part of its input, and produces a new hidden state as part of its output. The hidden state contains the information that the classifier needs to make a context-informed decision on the present sleep stage. This makes it a natural fit for performing real time inference, as one can perform context-informed inference on each new 30s epoch as it comes in, and does not need to observe the entire PSG at once.

### Accuracy

We compared the performance of the Greifswald Sleep Stage Classifier (GSSC) against three other recently developed, high-performance classifiers (Stephansen et al. 2018, U-Sleep, Perslev et al. 2021, YASA, Vallat and Walker, 2021) and found that it consistently outperformed Stephansen et al. (2018) and YASA (Vallat and Walker, 2021), and was at least at parity with U-Sleep (Perslev et al. 2021). This places GSSC at the current state of the art.

### Limitations

One disadvantage that GSSC has compared to YASA is speed. YASA can infer an entire night’s PSG in a few seconds, while GSSC requires somewhat more time, even with a GPU. With a CPU, inference time can go into the minutes for a full night - however this time can be significantly reduced by opting for a specific channel constellation rather than using the permutation consensus, likely at the cost of 1-2 percent accuracy. Researchers who have limited time and/or computing power, use conventional PSG setups and electrodes, and do not need the highest level of available accuracy might consider using YASA. Otherwise GSSC would be preferable for the reasons described above.

### Outlook

There are a number of things which should be improved in future versions of the GSSC. First, performance on the testing dataset over 20 training epochs did not converge, but rather oscillated continually (see figure SX). An average of the best three epochs produced excellent results on the validation sets, but it would nevertheless be preferable for the classifier to converge on a stable solution. Second, the classifier was trained on only 2652 PSGs from four datasets. This is somewhat less than YASA, a non-deep learning classifier (3,163 PSGs, seven datasets), and a small fraction of what was used for the deep learning-based U-Sleep (19,924 PSGs, 21 datasets). Training on more data may solve the non-convergence issue mentioned above, and also yield a non-trivial accuracy increase; on the other hand, performance at the current state of the art may already be near the intrinsic limits of how accurate sleep staging can be, given the relative indeterminacy of sleep staging criteria, and that machine/deep learning classifiers are trained on manually scored data, which are themselves quite variable (Rosenberg and Van Hout, 2013, Younes et al, 2016, Muto et al, 2018). Third, during prototyping, we found good performance on four-layer Resnets for the EEG channel, and one-layer Resnets for the EOG. This is unsurprising given how much more complex a brain signal is from an ocular muscle signal, but there may be space to more thoroughly fine-tune these layer numbers. Finally, the weights used to adjust the loss function for the severe imbalance of sleep stages could potentially be improved. We have simply adopted the ones reported for YASA, with a minimal change to the N1 weight (Vallat and Walker, 2021), and they have yielded excellent results, but some small adjustments could prove beneficial.

## Conclusion

The Greifswald Sleep Stage Classifier (GSSC) is free, open source, easy to install and use, offers state of the art accuracy, and performs well for all reasonable channel combinations, including only a single EOG channel. It is particularly well suited to real time inference (BCI, closed-loop stimulation). These features render the GSSC an excellent candidate for becoming a standard tool for polysomnographers.

## Availability

The Greifswald Sleep Stage Classifier is available at github.com/jshanna100/gssc/ under a GNU Aphero General Public License, version 3. This classifier is not certified as a medical device by any regulatory authority.

## Acknowledgements

This study was funded by a Sonderforschungsbereich project grant (327654276 – SFB 1315, B03) awarded to Agnes Flöel by the Deutsche Forschungsgemeinschaft. An NVIDIA TITAN V GPU used for this project was awarded to Jevri Hanna through the NVIDIA Academic Hardware Grant Program. Computing power provided by the High Performance Computing Cluster of the Free University Berlin also made a contribution to this research. We would like to thank Silke Wortha and Julia Ladenbauer for helpful consultations. Most figures were produced with the help of Matplotlib (Hunter, 2007) and Seaborn (Waksom, 2021). Neural network schematics were graphed with PlotNeuralNet v1.0.0 (Iqbal, 2018)

## Author contributions

Jevri Hanna conceived, designed, and tested the sleep stage classifier, and wrote the manuscript. Agnes Flöel provided supervision, funding, and wrote the manuscript.

## Competing interests

We declare no competing interests.

## Notes

https://github.com/jshanna100/gssc

### Competing Interest Statement

The authors have declared no competing interest.

