## Supplemental Info for "An accessible and versatile deep learning-based sleep stage classifier"

| | GSSC H | GSSC O | YASA H | YASA O | Perslev H | Perslev O | Stephansen H | Stephansen O |
|---|---|---|---|---|---|---|---|---|
| Acc | 88.8 +/- 4.8 | 88.1 +/- 5.8 | 86.2 +/- 7.3 | 83.2 +/- 6.0 | 89.5 +/- 2.2 | 86.1 +/- 8.1 | 85.9 +/- 5.8 | 84.9 +/- 9.3 |
| MCC | 82.5 +/- 6.8 | 81.8 +/- 10.3 | 78.0 +/- 10.2 | 75.6 +/- 9.9 | 84.3 +/- 3.5 | 79.4 +/- 11.1 | 80.1 +/- 8.3 | 77.5 +/- 11.7 |
| CK | 82.1 +/- 6.9 | 81.4 +/- 10.9 | 77.8 +/- 10.5 | 74.5 +/- 10.9 | 84.2 +/- 3.5 | 78.5 +/- 11.8 | 79.5 +/- 9.7 | 77.2 +/- 12.6 |
| F1 Macro | 81.8 +/- 7.9 | 79.9 +/- 10.6 | 78.7 +/- 11.1 | 74.0 +/- 11.6 | 83.3 +/- 4.5 | 79.4 +/- 10.4 | 78.2 +/- 5.5 | 69.8 +/- 15.9 |
| F1 Wake | 90.2 +/- 7.7 | 92.9 +/- 7.1 | 83.0 +/- 15.1 | 85.5 +/- 13.1 | 91.9 +/- 8.3 | 92.7 +/- 5.8 | 86.7 +/- 9.9 | 88.3 +/- 7.9 |
| F1 N1 | 56.2 +/- 21.1 | 45.7 +/- 22.7 | 50.0 +/- 12.3 | 40.5 +/- 18.9 | 60.6 +/- 13.1 | 53.3 +/- 16.7 | 51.8 +/- 20.2 | 41.3 +/- 18.5 |
| F1 N2 | 90.9 +/- 5.6 | 89.7 +/- 7.1 | 88.4 +/- 5.0 | 86.1 +/- 5.9 | 89.7 +/- 5.3 | 87.6 +/- 8.4 | 89.4 +/- 5.6 | 88.4 +/- 7.8 |
| F1 N3 | 88.1 +/- 11.9 | 81.4 +/- 29.7 | 88.6 +/- 13.7 | 75.9 +/- 23.5 | 85.2 +/- 13.8 | 76.4 +/- 33.8 | 75.3 +/- 23.5 | 67.3 +/- 69.6 |
| F1 REM | 94.3 +/- 4.6 | 93.3 +/- 5.1 | 92.7 +/- 10.2 | 89.2 +/- 10.5 | 94.1 +/- 5.0 | 93.6 +/- 6.5 | 91.5 +/- 5.8 | 90.3 +/- 15.7 |

**Table S1: Performance of the GSSC, YASA, U-Sleep (Perslev et al. 2021) and Stephansen et al. (2018) on the DREEM Health and Obstructive datasets.** Measures include Accuracy (Acc), Matthews Correlation Coefficient (MCC), Cohen's Kappa (CK), F1 Macro and F1 scores for each individual stage. These are depicted graphically in Figs. 3 and S4.

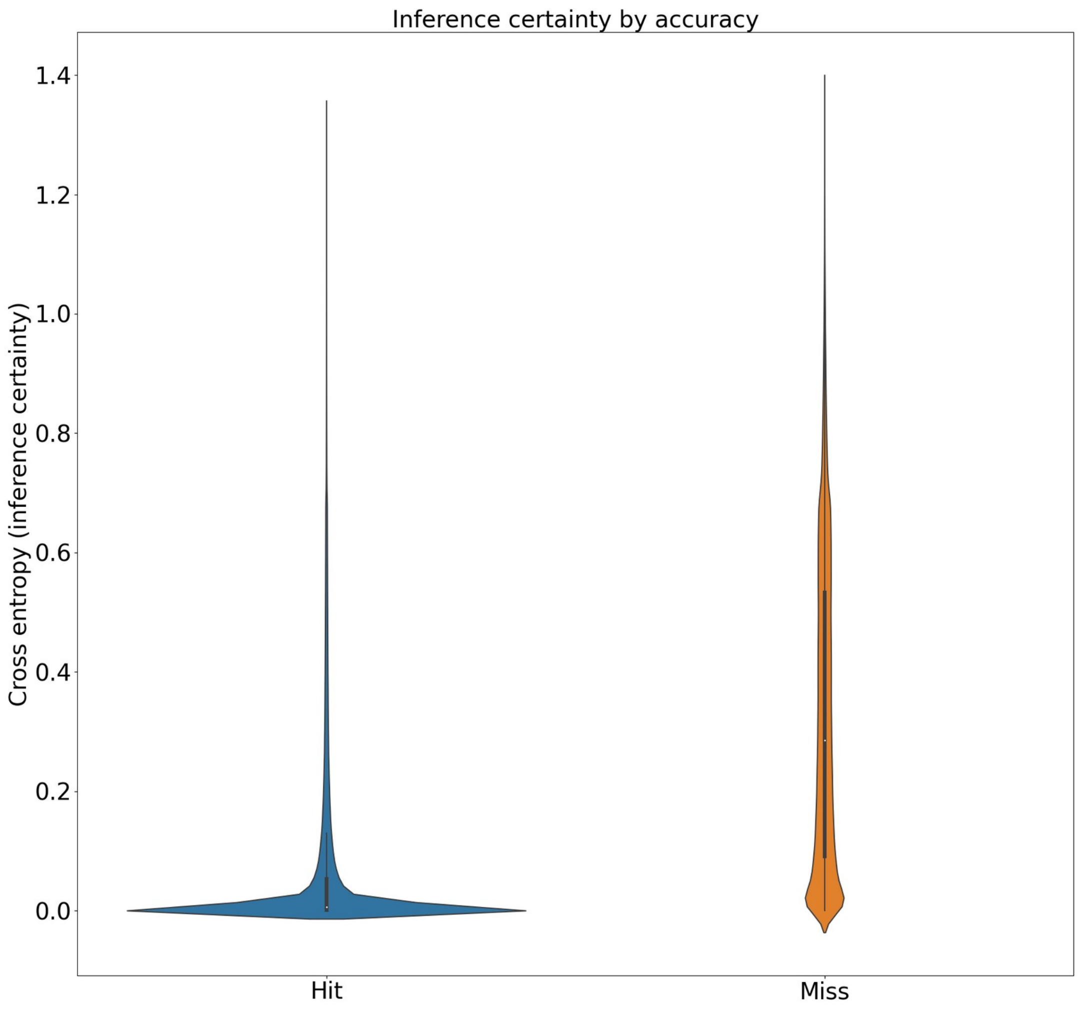


**Figure S1: Relationship of inference certainty and accuracy.** Violin plots show sross entropy loss of inferred sleep stage against the log softmax output of the classifier, which can be understood as an inverse index of the classifier's certainty about its inference. Accurate inferences ("Hit") tend to have much more certainty than inaccurate inferences ("Miss").

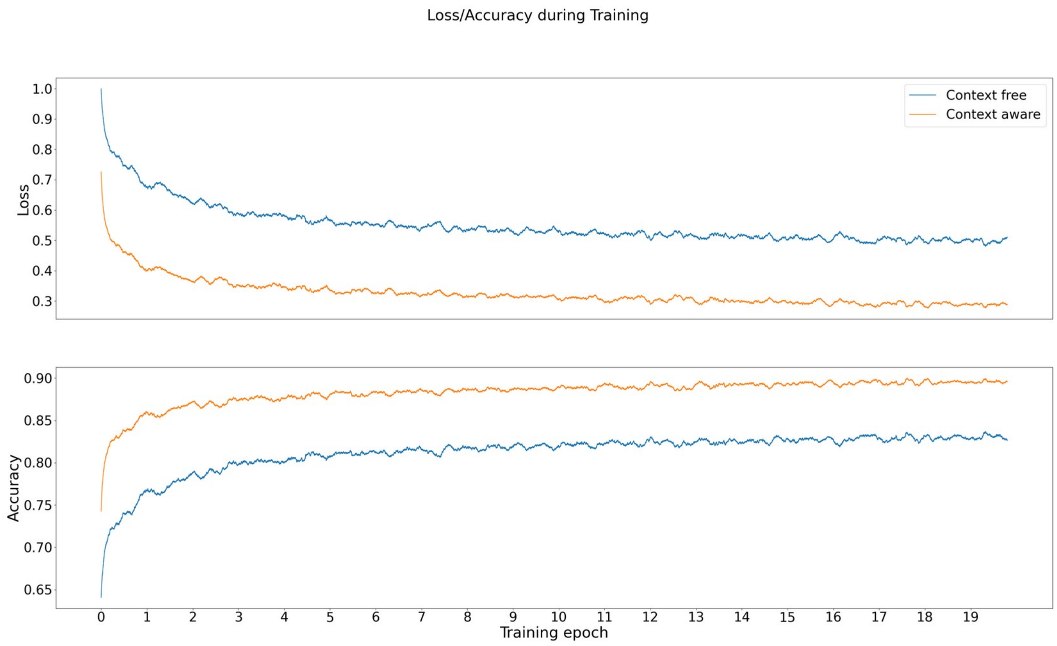


**Figure S2: Loss and accuracy across 20 training epochs.**

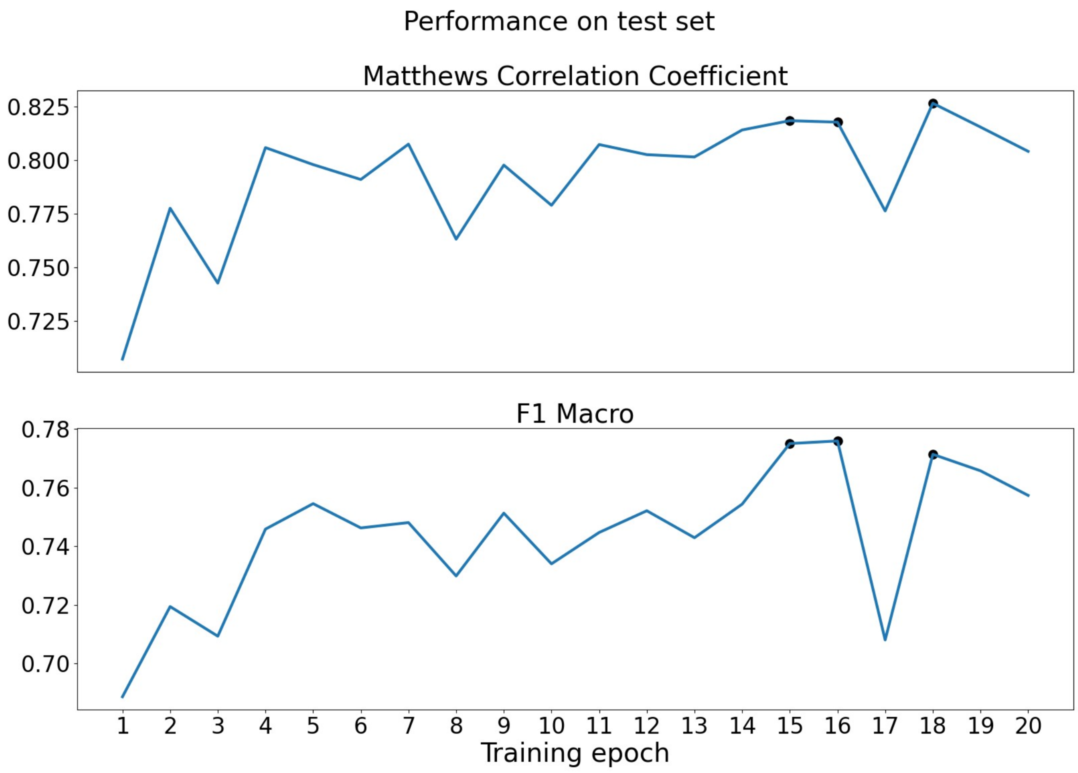


**Figure S3: Performance on the test set as assessed by MCC and F1 Macro.** The MCC is a good assessment of overall performance, while the F1 Macro requires good performance on all possible classes. Because the model did not converge on the test set, three points were selected that had high scores on both measures, and the network weights for these three points were averaged. This network was then applied to the validation set to produce the results reported in this paper.

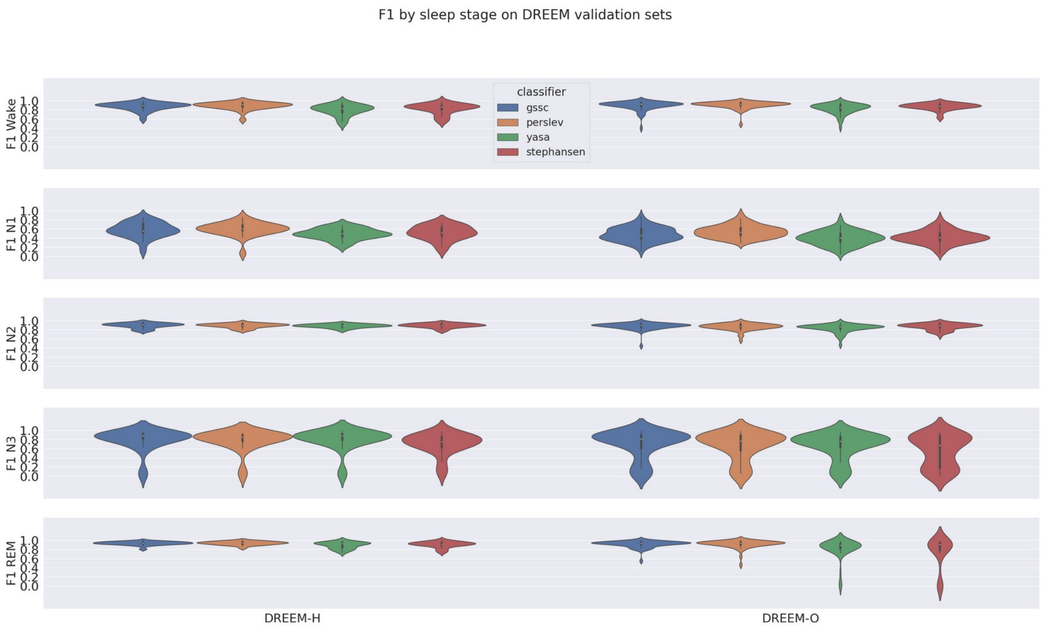


**Figure S4: F1 measures for four classifiers for the five possible sleep stages on the DREEM validation datasets.**
